# Tactile innervation densities across the whole body

**DOI:** 10.1101/2020.04.27.063263

**Authors:** Giulia Corniani, Hannes P. Saal

**Affiliations:** Active Touch Laboratory, Department of Psychology, University of Sheffield, Sheffield S1 2LT, UK; Sheffield Robotics, University of Sheffield, Sheffield S1 3JD, UK; INSIGNEO Institute for in silico Medicine, University of Sheffield, Sheffield S1 3JD, UK

**Keywords:** cutaneous afferent, mechanoreceptor, glabrous skin, hairy skin

## Abstract

The skin is our largest sensory organ and innervated by afferent fibers carrying tactile information to the spinal cord and onto the brain. The density with which different classes of tactile afferents innervate the skin is not constant but varies considerably across different body regions. However, precise estimates of innervation density are only available for some body parts, such as the hands, and estimates of the total number of tactile afferent fibers are inconsistent and incomplete. Here we reconcile different estimates and provide plausible ranges and best estimates for the number of different tactile fiber types innervating different regions of the skin, using evidence from dorsal root fiber counts, microneurography, histology, and psychophysics. We estimate that the skin across the whole body is innervated by approximately 230,000 tactile afferent fibers (plausible range: 200,000-270,000). 15% innervate the palmar skin of both hands and 19% the region surrounding the face and lips. Around 60% of all tactile fibers are slowly-adapting, while the rest are fast-adapting. Innervation density correlates well with psychophysical spatial acuity across different body regions, and additionally, on hairy skin, with hair follicle density. Innervation density is also weakly correlated with the size of the cortical somatotopic representation, but cannot fully account for the magnification of the hands and the face.

## Introduction

Understanding sensory processing requires an appreciation of the nature of sensory inputs. Careful study of the visual system has revealed that about 100 million photoreceptors in each retina convert light into electrical impulses, which are relayed through roughly 1 million retinal ganglion cells in the optic nerve (1). In audition, about 12,000 hair cells in each cochlea pass on auditory information to the brain (2). The sense of touch puts to use our largest sensory organ, the skin, which is innervated throughout by cutaneous fibers signaling light touch, temperature, and pain. Despite its importance for manipulation (3), movement (4), our sense of body ownership (5), and affection (6), we know little about the number and distribution of cutaneous fibers innervating different skin regions across the body. Estimates of tactile fiber innervation in the current literature are few, often incomplete and inconsistent, and range from a total innervation of around 45,000 fibers (7) into the millions (8). Most textbooks do not even venture a guess (9–12). Reliable estimates exist only for a few regions of glabrous skin. The gold standard is a study by Johansson and Vallbo (13) that estimated that around 18,000 myelinated tactile fibers innervate the palmar surface of each hand.

Various techniques can be employed for counting fibers, but individually they all suffer from problems, which explains the discrepancy in estimates. Histological examination can provide estimates for the number of fibers in the peripheral nerves, but cannot distinguish between afferent and efferent fibers. Furthermore, peripheral nerves carry many types of sensory fibers other than tactile ones, for example, proprioceptive fibers or those innervating internal organs, such as the bladder. Immunohistochemistry of samples taken by skin biopsies allow receptor and fiber counts, but the regions covered are necessarily very small, and innervation of the skin is not uniform. Individual tactile fibers often branch and innervate tens of receptors, and estimates of branching and convergence factors differ widely. Another approach estimates innervation density from psychophysically determined two-point discrimination thresholds. Here, the idea is that higher innervation density enables improved spatial localization, so finer spatial discrimination should be associated with higher fiber count. However, such estimates often ignore that discrimination thresholds can improve with training (14) and likely rely predominantly on only one of the multiple different afferent classes that innervate the skin (15). The general problem is that none of the methods described above can be used to extrapolate between glabrous and hairy skin without taking into account the different composition in the types of tactile afferent fibers. Finally, much valuable insight into the prevalence of different fiber types also comes from microneurography, a technique for obtaining electrophysiological recordings from individual fibers in human nerves. However, this technique has mostly been applied to fibers terminating in the hand, the foot, or the face. A handful of studies has investigated the hairy skin of the hand dorsum and arm, but data from other body regions is sorely lacking. Data from animal models, specifically primates, can also provide valuable input, however stark differences in innervation density have been observed across different primate species (16), so such data can only be used with caution.

Here, we combine published evidence from all available measures (histology of the nerve and the skin, microneurography, psychophysics) to estimate plausible ranges for innervation densities of Aβ myelinated tactile afferent fibers covering all skin regions of the body. We estimate that the skin is innervated by approximately 230,000 tactile afferent fibers (plausible range: 200,000-270,000) in total. The hands and the face are the most highly innervated skin regions, as might be expected from the exaggerated cortical representation of these body parts (17). While we believe our estimates are robust, more basic work remains to be done, especially concerning the innervation of hairy skin.

## Innervation of the skin

The sensory innervation of the skin has been extensively covered in reviews (6, 18–20) and textbooks (10), so we will only provide a brief overview here.

The main tactile fibers underlying discriminative touch are myelinated Aβ fibers. Some tactile information is also carried by slow, unmyelinated C fibers (e.g., C-tactile fibers), which are thought to be important for affective touch (21). Due to a paucity of research on these fibers and their extensive branching within the skin, it is hard to establish reliable estimates on their prevalence. Moreover, unmyelinated fibers are most commonly associated with sensing pain and temperature and cannot be distinguished histologically from their tactile cousins. In the following, we will, therefore, focus on Aβ fibers exclusively.

Two major afferent classes exist, which are distinguished by their electrophysiological response properties: fast-adapting (FA) fibers (also called RA: rapidly-adapting or QA: quickly adapting in the literature) that respond exclusively to dynamic stimuli, that is when the skin is in motion; and slowly-adapting (SA) fibers, which, in addition to dynamic responsiveness, also respond to sustained static skin deformation and stretch. Both classes can be further subdivided into type I afferents, which are more numerous and terminate close to the surface of the skin, and type II afferents, which end in deeper skin layers. The presence and prevalence of different afferent classes differ in glabrous (non-hairy) skin as found on the palm, sole, and the lips, as compared to hairy skin, which covers the rest of the body.

Different afferent classes are associated with different mechanoreceptors. However, some of these associations are still debated and might not apply to all skin types. In the following, we will note associations that have been made in the literature, but our estimates will apply only to electro-physiologically characterized afferent types, and we make no claim regarding their associated mechanoreceptors. We will report innervation densities as units/cm^2^, where we take a unit as the structure composed of an afferent fiber and all the mechanoreceptors (if any) innervated by it.

## Glabrous skin of the hand

Unlike other body regions, the glabrous skin of the hand and its tactile afferent innervation have been extensively studied, owing to its importance in grasping and manipulation. The number of tactile afferent fibers in the glabrous skin of the hand of young adults is estimated as between 17,000 and 19,000 (13). There are slightly more fast-adapting fibers (56%) than slowly-adapting ones (44%), a common feature of glabrous skin (see section on the foot sole).

Four major afferent types have been identified in the palmar skin of the hand: fast-adapting type I (FAI) fibers that innervate Meissner corpuscles; slowly-adapting type I (SAI) fibers that innervate Merkel cells; slowly-adapting type II (SAII) fibers that innervate Ruffini corpuscles; and fast-adapting type II (FAII) fibers that innervate Pacinian corpuscles.

43% of tactile afferent fibers or around 7,740 fibers are fast adapting type I fibers (FAI). FAI afferents are densely packed in the human fingertip with 141 units/cm^2^ at its distal end. The density decreases in the proximal direction and only 25 units/cm^2^ are present in the palm (see Figure 1). The end organs of FAI fibers are Meissner corpuscles (MCs). Each Meissner corpuscle is innervated by one or two Aβ myelinated FA fibers (22), and a single FA fiber typically branches to innervate several MCs (23). In the human fingertip, up to 50 MCs/mm^2^ can be found (16). Meissner corpuscle density in the palm is considerably lower with 5 MCs/mm^2^ at the thenar eminence (24).

**Fig. 1.**
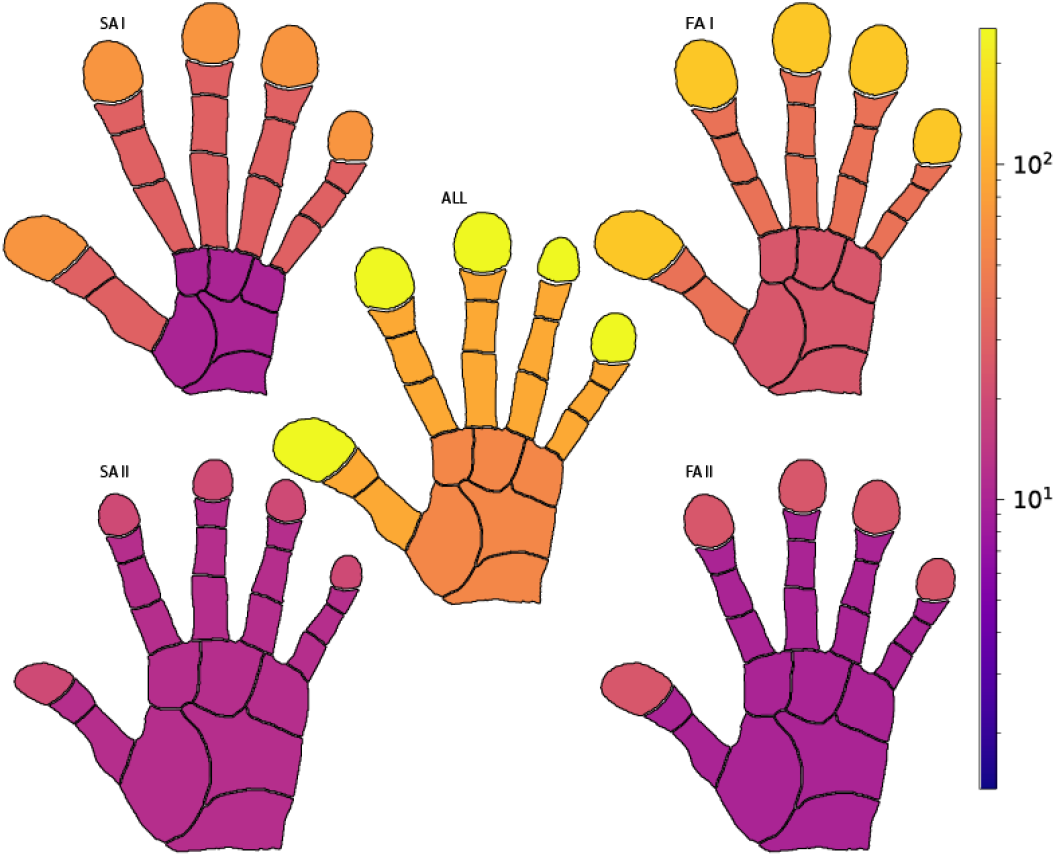
Innervation of the palmar surface of the human hand for different afferent classes. Colors denote different innervation density (units/cm^2^). Each hand area is scaled by its innervation density to reveal the hand ‘homunculus’. Both SA I and FA I fibers are densely packed in the distal ends of the fingertips and much less so in the palm, while the two other afferent classes are more evenly spread throughout the hand and exhibit much lower innervation density overall.

25% of tactile afferent fibers or around 4,500 fibers in the palmar region of a single hand are classed as slowly adapting type I fibers (SAI). SAI fibers are densely concentrated in the fingertips at around 70 units/cm^2^ at its distal end, and less so in the more proximal area of the hand with 46 units/cm^2^ in the middle phalanx and 10 units/cm^2^ in the palm (see 1). SAI fibers repeatedly branch and innverate Merkel cell neurite complexes, which form clusters within the skin. In the fingerpad of normal adults, up to 10,000 Merkel cells/cm^2^ can be found, but not all of them appear to serve mechanoreceptive functions or are connected to nerve fibers (25).

19% of tactile afferent fibers or around 3,400 fibers are classed as slowly adapting type II fibers (SAII). These are uniformly distributed across the glabrous skin area of the hand at an innervation density of approximately 12 units/cm^2^. However, there is some evidence for increased density at the skin/nail border on the fingertips (13, 26). SAII fibers innervate Ruffini corpuscles (27), but it is unclear whether all SAII-like responses originate from Ruffini corpuscles. Where they do, a one-to-one mapping between fibers and corpuscles is assumed (28).

Finally, 13% of tactile afferent fibers or around 2,300 fibers are fast-adapting type II (FAII). FAII fibers target Pacinian corpuscles. There is a one-to-one correspondence between FAII fibers and Pacinian corpuscles, which means that counting corpuscles can serve as a direct estimate of FAII fiber count. The distribution of this mechanoreceptor type is almost uniform across the hand surface, with an estimated total of 800 corpuscles in the palm and 350 corpuscles in each finger (13). Pacinian corpuscles are relatively large and can, therefore, easily be identified in dissections, at least in principle. They appear to form clusters close to the digital nerves and their branches. Examinations of the hands of human cadavers have counted around 300 corpuscles per hand (29), much fewer than estimated by Johansson and Vallbo (13). However, it should be noted that these studies were done on elderly individuals, who might have experienced a loss of corpuscles during aging. In support of this idea, Cauna and Mannan (30) counted Pacinian corpuscles in the radial half of a foetal index finger and found 178 in total, in almost perfect agreement with Johansson’s estimates. More recently, it has been shown that Pacinian corpuscles can be resolved using high-field MRI (31, 32), but this technique has not yet been used to establish precise counts.

Receptive fields of the type I fibers on the glabrous skin of the hand are small, circular, and well-defined with a mean area of 13 mm^2^ for the FAI and 11 mm^2^ for the SAI fibers. Receptive fields of type II fibers are larger with diffuse borders and a mean area of 101 mm^2^ for FAII and 59 mm^2^ for SAII fibers (33).

## Glabrous skin of the foot sole

Somatosensory feedback from the lower limb, and in particular from the foot sole, plays an important role in controlling balance, posture, and gait (34). The foot sole is covered with glabrous skin and innervated by the same four classes of tactile afferents as the hand (SAI, SAII, FAI, FAII).

We estimate the total number of plantar cutaneous tactile afferent fibers for a single foot sole to be around 4,000, divided as follows: 17% (~680) SAI fibers, 20% (~800) SAII fibers, 51% (~2,040) FAI fibers, and 12% (~480) FAII fibers. These numbers are higher than an earlier estimate provided by Strzalkowski et al. (35) (see Methods for details regarding our estimation methodology). Like the hand, the foot sole contains more fast-adapting (62%) than slowly-adapting fibers (38%). The distribution of cutaneous afferents is not uniform across the foot sole for type I afferents (2). The overall highest innervation density is found in the toes (48 units/cm^2^), followed by the lateral arch (31 units/cm^2^), the lateral metatarsals (29.7 units/cm^2^) and the heel (15.7 units/cm^2^). The lowest innervation density was in the medial metatarsals (11.3 units/cm^2^). FA afferents are considerably more dense in the toes (24.5 units/cm^2^) than in the metatarsal/arch (9.1 units/cm^2^) and in the heel (8 units/cm^2^). A similar distribution is observed for SAI afferents. Similar to the hand, SAII and FAII fibers are almost uniformly distributed across the different areas of the foot sole skin (Figure 2). In electrophysiological recordings, fewer tactile afferents have been found terminating in the great toe as would be expected given its size, and its innervation thus appears lower than that of the neighboring toes; whether this discrepancy reflects a statistical artefact or a genuine difference remains to be seen.

**Fig. 2.**
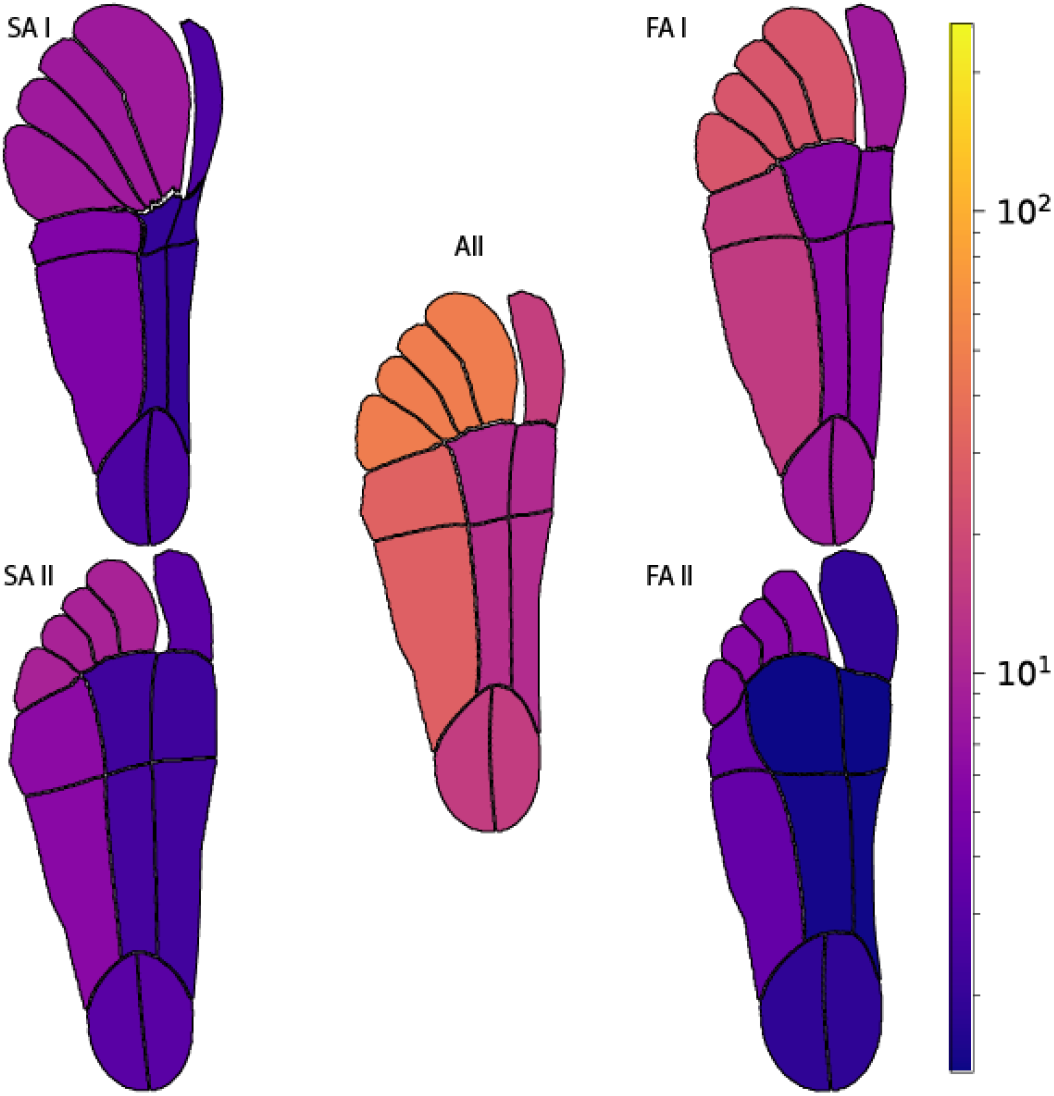
Innervation densities (units/cm^2^) for different afferent classes across the plantar surface of the human foot. Compared to the hand, the foot sole is much less densely innervated but displays a similar proximal-distal gradient for type I afferents. Additionally, a lateral gradient is evident with dense innervation of the lateral arch

The size of the receptive fields varies considerably for the different tactile fibers and across different foot areas with a mean value of 76 mm^2^ for the SAI fibers, 248 mm^2^ for SAII fibers, 81 mm^2^ for FAI fibers, and 873 mm^2^ for FAII fibers. In general, larger receptive fields are reported in the middle metatarsal and heel, and smaller receptive fields are located in the toes (35). Receptive fields on the foot sole are thus several times larger than those measured in the hand, perhaps owing to the less dense innervation of this skin region and the different mechanical properties of the skin of the foot sole.

## Face

The face is densely innervated by cutaneous fibers, especially the region around the mouth and lips, as well as the inside the oral cavity and the tongue, highlighting the essential sensory contribution to mastication and other eating-related behaviors. We estimate that around 43,000-46,000 tactile afferents innervate the hairy facial skin and the lips, excluding the oral cavity, which is likely to be innervated by around 16,000-19,000 fibers (see Methods for details).

Both slowly and fast adapting tactile afferents are found in the hairy skin of the face and the red zone of the lip and mucosa. Slowly adapting afferents appear more abundant than fast adapting ones both in the facial skin and in the labial oral mucosa (36, 37), with around 65% SA, resulting in 29,000 fibers, and 35% FA, resulting in 15,500 fibers. However, this breakdown is extrapolated from relatively small samples, so should be treated with caution. Notably, no FAII afferents have been reported in the literature, and vibrotactile thresholds on the face show no characteristic Pacinian sensitivity around 200 Hz (38), so this afferent class appears absent on the face, while present in all other body regions. It should be noted, however, that extremely light stimulation that is usually associated with FAII afferents, such as a gentle puff of air, is readily experienced even on facial skin. It is therefore conceivable that FAII are present yet rare in facial skin. There is a dearth of descriptions of Pacinian corpuscles in hairy skin generally, suggesting that these might generally be located deeper in the skin and their numbers might be low. Innervation density is not uniform across the face: we estimate an innervation density of 48 units/cm^2^ for the forehead, eyes, and nose (V1), 66 units/cm^2^ for the central part of the face (V2) and 83 units/cm^2^ for the lower lip, the chin, the jaw and an area around the ears (V3). Locally, some regions like the area immediately surrounding the mouth and the lips are likely to exhibit much higher innervation densities.

The size of the receptive fields varies for the different tactile afferent fibers with a mean value of 4 mm^2^ for the SAI fibers, 6 mm^2^ for SAII fibers and 6 mm^2^ for FA fibers (36). Most receptive fields have a circular or oval well-demarcated area of high and relatively uniform sensitivity (37). The highest concentration and smallest size of the receptive fields are measured around the corner of the mouth and in the upper lip. The psychophysical and receptive field properties observed in these areas, including the tactile acuity, are similar to those found in the human fingertip (39), suggesting a similarly high innervation density.

## Hairy skin

The sensory innervation of human hairy skin (other than facial) has only been scarcely investigated, and microneurography studies have restricted themselves to the hand dorsum (40? –42), the arm (43), the leg, and the foot dorsum (44). Hairy skin is innervated by afferent classes with similar response characteristics as found in glabrous skin, though specific end organs might differ. As in all other types of skin, SAI afferents are present and innervate Merkel cells, which in hairy skin are organised into touch domes, as compared to the cell neurite complexes found in glabrous skin. Similarly, SAII afferents have been identified electrophysiologically, though it is unclear whether they always connect to Ruffini-like corpuscles, as is thought to be the case in the hand (45). Afferents with response properties similar to FAI afferents are frequently observed, but unlike glabrous skin, hairy skin does not contain Meissner corpuscles. Instead, FAI fibers branch and terminate in close proximity to hair follicles (‘hair units’). Each FA fiber is estimated to innervate around 25 individual hairs in the forearm (43). The microneurography literature also describes FAI field fibers, which appear unconnected to hairs, but otherwise exhibit typical response properties similar to FAI fibers. The presence of FAII afferents has been demonstrated both in electrophysiological recordings (43) and psychophysically (46), though Pacinian corpuscles appear extremely rare in hairy skin. Overall, hairy skin is innervated by more SA fibers (65%) than FA ones, with estimates consistent across different studies (41, 43), though data is limited to the arm.

Based on fiber counts and estimates of axon diameter distribution in the dorsal roots of the spinal cord (see Methods), we estimate that around 145,000 Aβ fibers (range: 110,000-180,000) innervate the hairy skin of humans (excluding the face). Innervation is most dense in the back of the head and neck area with around 22 units/cm^2^ and almost uniform in the rest of the body, with 9 units/cm^2^ covering the skin of the arms and the hand dorsum, 10 units/cm^2^ in the chest and abdomen, 10 units/cm^2^ on the back, and 9 units/cm^2^ for the legs.

Hair units terminate on hair follicles in hairy skin, suggesting a relationship between hair follicle density and FA afferent innervation. Hair follicle density is not uniform across the adult body but instead varies by more than an order of magnitude across different body regions (47). If the number of hair follicles innervated by a single afferent was relatively constant across different areas, one would, therefore, expect a strong correlation between our estimates of FAI innervation density and hair follicle density. Indeed, we found a strong correlation of *r* = 0.962 (*p* = 0.002) between both quantities (Figure 3). Our estimates imply that each FAI hair fiber innervates, on average, 24 hair follicles (range: 15-42), in strong agreement with earlier estimates for the forearm (43).

**Fig. 3.**
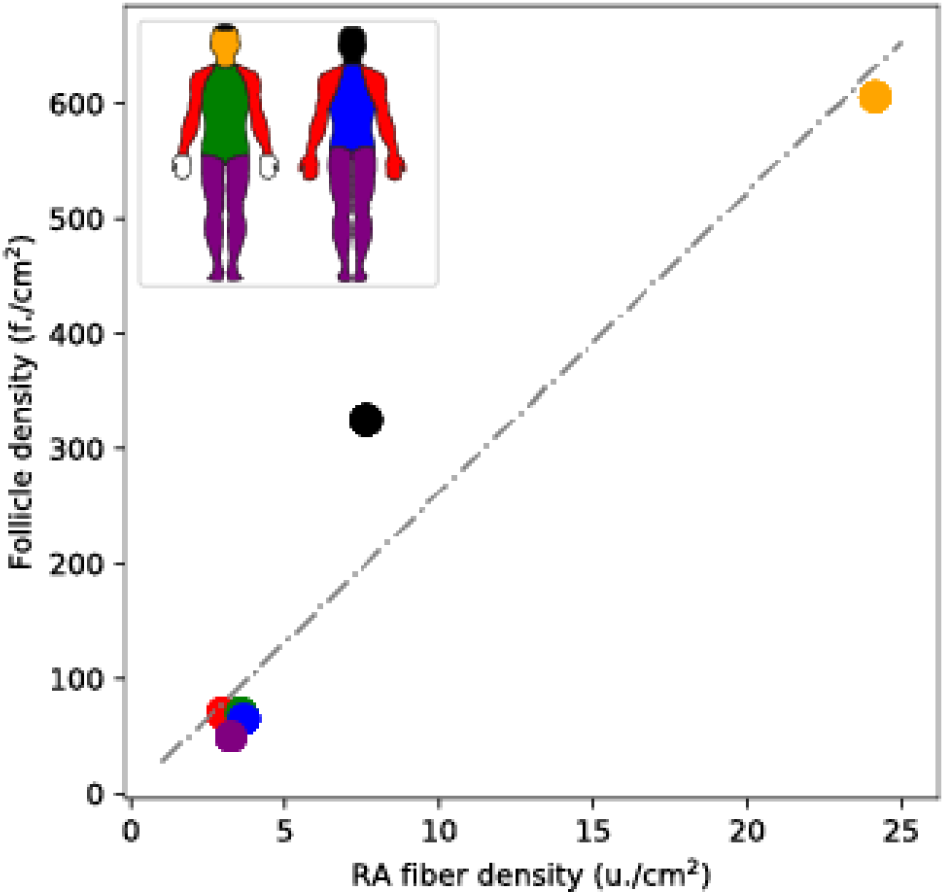
Estimated fiber density for FA hair cells on different skin regions versus average hair follicle density for the same skin regions.

## Whole-body innervation

Across the whole body, the palmar skin of the hands and the perioral region of the face are the most densely innervated regions. Relatively high innervation can also be found in some sections of the foot, such as the toes, while the hairy skin of the arms and legs are the least densely innervated, closely followed by the trunk (see Figures 4 and 5, and Table 1).

**Fig. 4.**
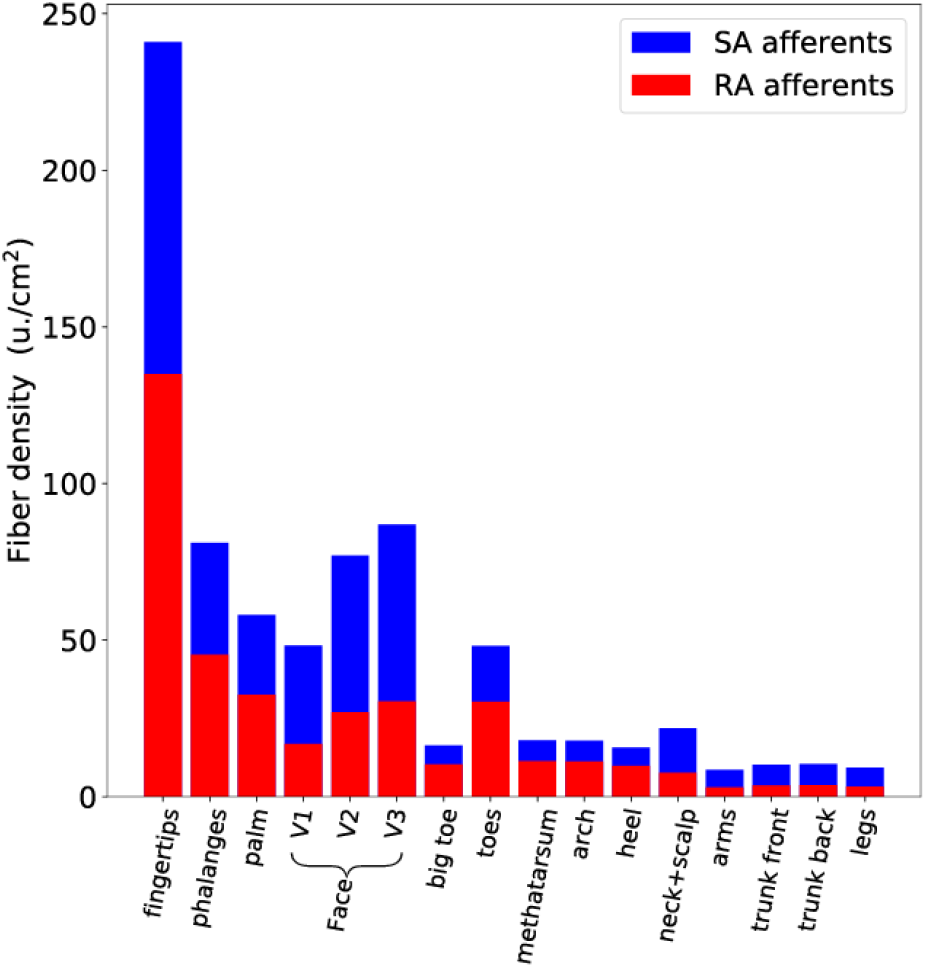
Total tactile innervation density for fast-adapting (red) and slowly-adapting (blue) afferents (including both type I and type II afferents), for different skin regions across the whole body. The hand and face are the most densely innervated regions. The ratio of fast and slowly adapting fibers is not constant but varies with skin region.

**Fig. 5.**
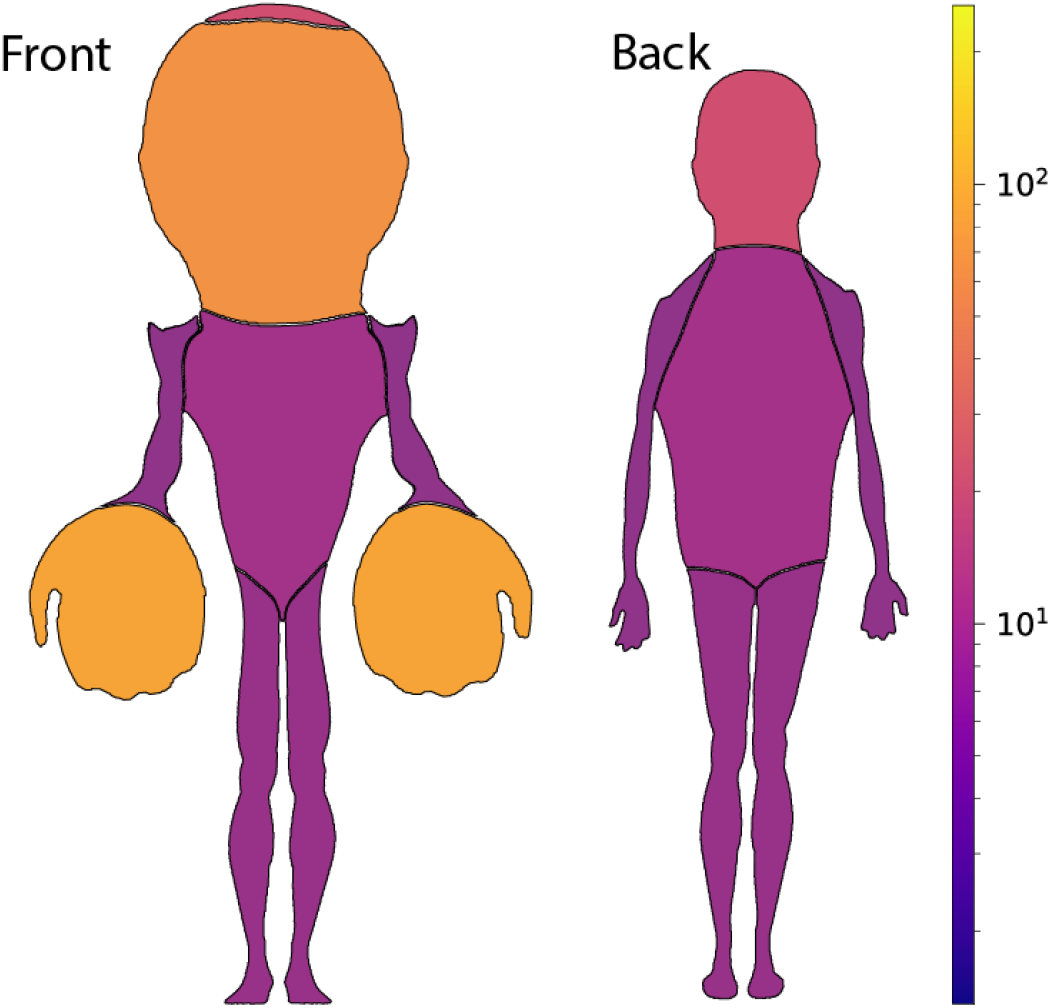
Whole-body tactile innervation densities. Colors and scale of each body area denote different innervation density (units/cm^2^). The hand and the face are the most densely innervated body parts.

**Table 1.**
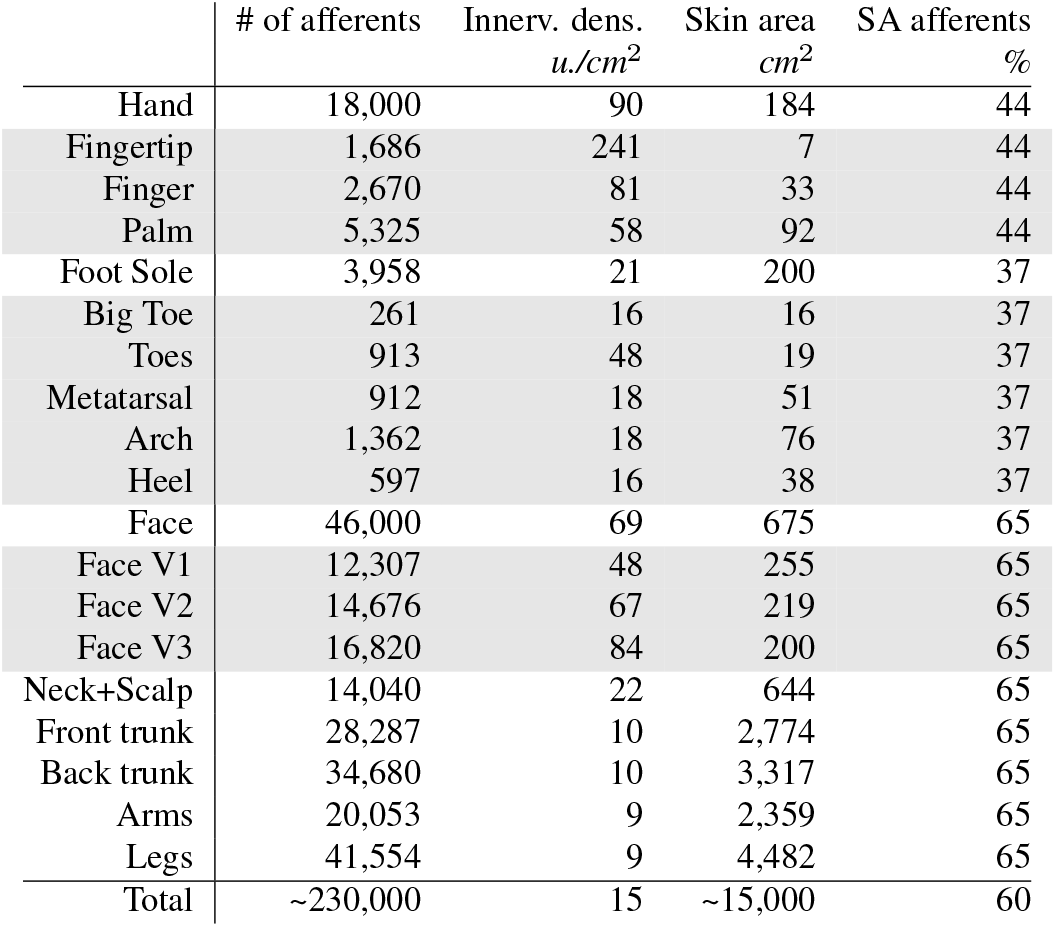
Estimated number of afferents, innervation density, skin area, and proportion of slowly-adapting afferents for different body regions. Shaded rows indicate sub-regions of larger body parts.

|  | # of afferents | Innerv. dens.<br>u./cm<sup>2</sup> | Skin area<br>cm<sup>2</sup> | SA afferents<br>% |
| --- | --- | --- | --- | --- |
| Hand | 18,000 | 90 | 184 | 44 |
| Fingertip | 1,686 | 241 | 7 | 44 |
| Finger | 2,670 | 81 | 33 | 44 |
| Palm | 5,325 | 58 | 92 | 44 |
| Foot Sole | 3,958 | 21 | 200 | 37 |
| Big Toe | 261 | 16 | 16 | 37 |
| Toes | 913 | 48 | 19 | 37 |
| Metatarsal | 912 | 18 | 51 | 37 |
| Arch | 1,362 | 18 | 76 | 37 |
| Heel | 597 | 16 | 38 | 37 |
| Face | 46,000 | 69 | 675 | 65 |
| Face V1 | 12,307 | 48 | 255 | 65 |
| Face V2 | 14,676 | 67 | 219 | 65 |
| Face V3 | 16,820 | 84 | 200 | 65 |
| Neck+Scalp | 14,040 | 22 | 644 | 65 |
| Front trunk | 28,287 | 10 | 2,774 | 65 |
| Back trunk | 34,680 | 10 | 3,317 | 65 |
| Arms | 20,053 | 9 | 2,359 | 65 |
| Legs | 41,554 | 9 | 4,482 | 65 |
| Total | ~230,000 | 15 | ~15,000 | 60 |

### Innervation density and tactile acuity

Innervation density limits the spatial resolution with which tactile features can be resolved on the skin: lower innervation results in a larger spacing between receptors and implies that two tactile stimuli need to be further apart to be discriminated. One might, therefore, expect a strong correlation between receptor spacing and perceptual tactile acuity as determined in psychophysical experiments. Previous work suggests that spatial acuity is largely driven by SAI afferents (15), which possess the smallest receptive fields and, therefore, the highest spatial resolution. Indeed, a close relationship between SAI receptor spacing and tactile acuity has been established across the different regions of the hand (48, 49). Following this line of research, we used psychophysical two-point discrimination thresholds obtained from different regions across the whole body (50, 51) and correlated these values with estimated SAI receptor spacing. We found a strong relationship between these two variables (*r* = 0.93, *p* < 0.001, see Fig. 6). As prior research has shown, tactile acuity is not fixed but improves with training; while the eventual plateau performance is likely determined by innervation density, typical performance might not be (15). Nevertheless, differences in innervation density across the whole body appear large enough to yield a reliable correlation with psychophysical thresholds.

**Fig. 6.**
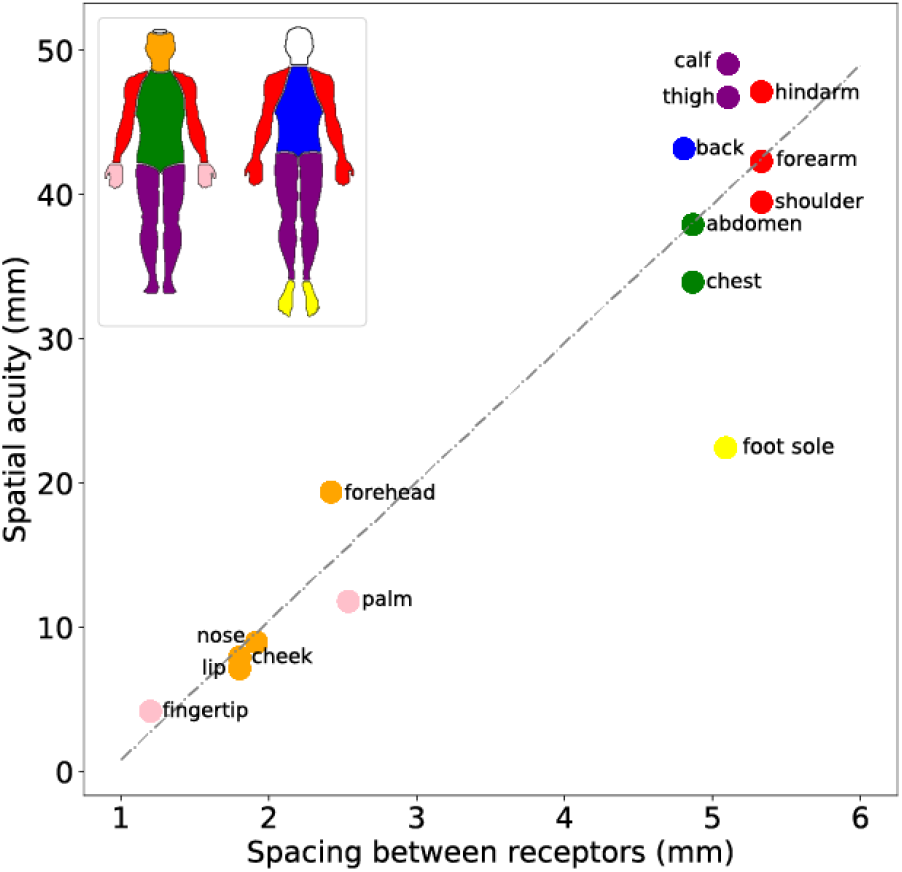
Estimated spacing between SAI termination sites versus perceptual tactile acuity as assessed by two-point discrimination tasks for different body regions (50). There is a strong relationship between a body region’s tactile innervation and our ability to spatially discriminate tactile stimuli.

### Innervation density and the cortical homunculus

As first investigated by Wilder Penfield (17, 52), body regions are mapped somatotopically onto the primary somatosensory cortex (S1), with nearby regions on the body generally represented by nearby patches in the cortex. However, the size of individual body regions in the cortex is not proportional to that anatomical region’s skin surface area. For example, the area devoted to the thumb in S1 is as large as the area devoted to the entire forearm (10). These findings have led to the famous homunculus, in which body parts are scaled by the size of their cortical representation, and which displays enlarged hands, face, and tongue. In how far cortical magnification is driven purely by innervation density, or whether usage effects such as increased contact with some body parts over others also play a part, has been debated in the literature. Many studies and textbooks argue for a close correlation between innervation density and cortical magnification (11, 53), though quantitative evidence is lacking. To test this idea directly, we took estimates of cortical magnification from the literature (52, 54), and compared these with the innervation density estimates described above. We found a low, but positive correlation between a region’s peripheral tactile fiber count and its representation in cortex (*r* = 0.10, *p* = 0.84, Figure 7), when assessed as the length of the coronal section onto which that body part is mapped. Crucially, some regions exhibited much larger cortical magnification than would be expected from their peripheral innervation alone. This included the heavily enlarged cortical areas containing the hand and face representation. Thus, it appears that these body parts are further magnified cortically, perhaps reflecting the fact that these are more likely to receive tactile stimulation or that they are especially behaviourally relevant. Interestingly, this cortical magnification of regions with already high innervation mirrors the visual system, where the fovea is further magnified cortically beyond its already much higher density of cone photoreceptors (55).

**Fig. 7.**
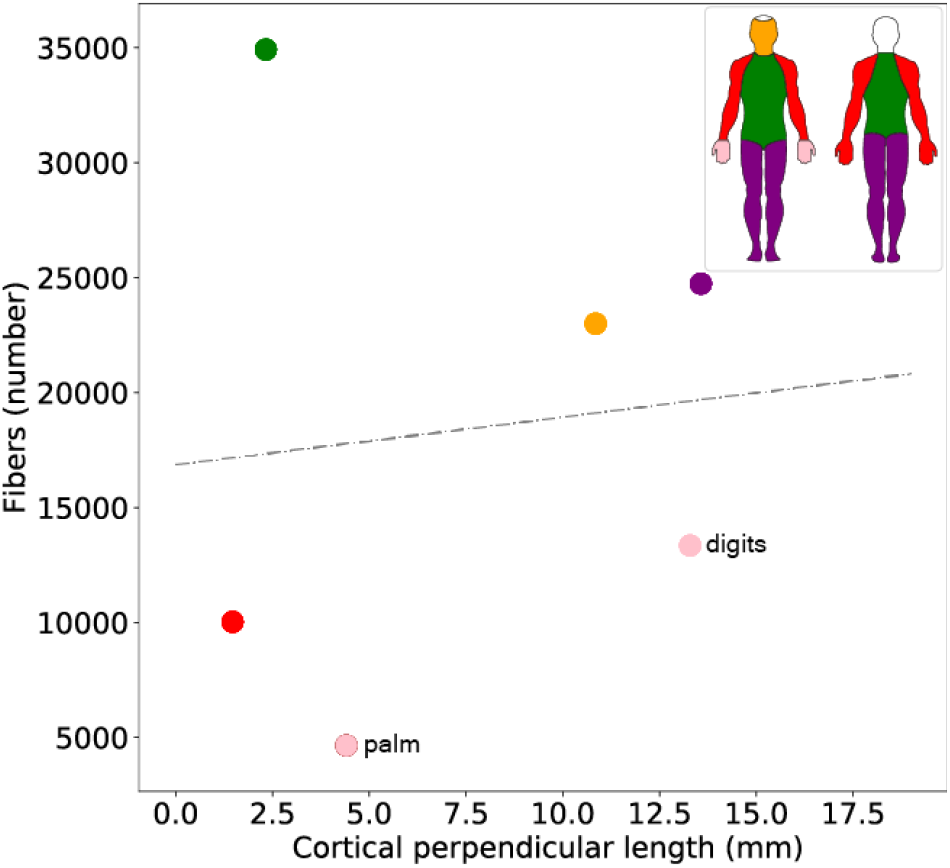
Size of cortical somatosensory representation for different body parts versus estimates for the total number of tactile fibers innervating that region. Innervation alone cannot explain cortical representation. Numbers refer to a single brain hemisphere.

### Tactile innervation over the lifespan

Our estimates of innervation density are based on data from a wide range of ages, but we have tried, as much as possible, to focus on young adults. It has been widely demonstrated that tactile sensibility declines with age, as evidenced by increased sensory thresholds (56–59) and decreased spatial acuity (60, 61). This decline might be partially explained by age-related mechanical changes of the skin itself, such as in stiffness or moisture levels, but neural degeneration through changes in myelination, receptor and fiber loss are likely to play a major part.

Focusing on the loss of tactile afferents specifically, a substantial decrease in the number of myelinated fibers in the spinal cord dorsal root from early middle age onwards has been well documented (see 62, for an overview of the literature). After a significant increase in the number of myelinated fiber in the first life decade, a gradual loss of fibers occurs throughout the lifespan from the third decade onwards, with an approximate loss-rate of 5-8% per decade (63, 64). Therefore, the proposed estimate of 230,000 tactile afferents in the whole body of a young adult should be reduced to about 160,000 tactile afferents for people over 80 years old.

## Methods

Innervation densities and fiber counts have been estimated previously for the hand and the foot sole, but not for other body regions. Here, we combined evidence from dorsal root fiber counts, hair density, and psychophysical thresholds to fill these gaps, as well as re-evaluate previous estimates.

### Hand

For the palmar surface of the hand we follow the original estimates by Johansson and Vallbo (13), which agree well with later histological analyses: a count of myelinated fibers at both the metacarpophalangeal (MCP) joint, covering all fibers innervating a given finger, and at the terminal trifurcation, covering innervation of the fingertips only, yielded 2,100-4,800 fibers per finger and roughly 1,900-2,600 per fingertip (65). Assuming that around 45% of these fibers are tactile afferents in the Aβ range, similar to the proportion that has been estimated at the wrist (66), yields 1,000-2,200 tactile afferents per finger and roughly 800-1,200 per fingertip. These numbers agree remarkably well with Johansson and Vallbo’s estimates of 2,500 for the whole finger and 1,000 for the fingertip of the index finger (13). Psychophysical measurements also suggest that innervation density decreases dramatically between the fingertip and the palm, and SAI receptor spacing as calculated from Johansson and Vallbo’s estimates correlates highly with spatial acuity across different regions of the hand (48, 49).

### Foot sole

A recent study estimated that around 1,700 tactile afferents innervate the plantar surface of a single foot (35). Our estimates suggest that the actual innervation is likely higher, by a factor of approximately two, based on several observations. First, the original estimate was based on a study demonstrating a ratio of roughly 10:1 in myelinated fibers between the hand and the foot (65). However, fiber counts were only taken at the metacarpophalangeal and metatarsophalangeal joints, respectively, and because innervation density gradients are steeper on the hand than on the foot, this does not imply a ratio of 10:1 in the total fiber count. Instead, a ratio of 4-5:1 appears more realistic. Second, tactile acuity is higher on the foot sole compared to the foot dorsum or other regions on the leg (51), suggesting a higher innervation density in this region, in line with our estimates for hairy skin (see methodology below). Taken together, a total tactile innervation of 4,000 afferents per foot sole appears most likely. To arrive at updated estimates for different regions of the foot sole, we took the total number of tactile afferents as estimated above and distributed them across the foot sole according to the relative densities established in Strzalkowski et al. (35).

### Face

The sensory innervation of the face is supplied by the sensory root of the trigeminal nerve or fifth cranial nerve. In this root, the total number of fibers is estimated at 170,000, and approximately 62,000 of these are myelinated and fall within the diameter range of Aβ fibers (67). The trigeminal nerve branches into three major divisions which supply different areas of the face; the ophthalmic branch, or V1, innervates the upper part of the face, covering approximately 38% of the facial skin; the maxillary branch, or V2, innervates the mid-third of the face, including part of the nose and down to the upper lip, corresponding to approximately 32% of the total facial area; finally, the mandibular branch, or V3, innervates the lower part of the face and the area around the ears, and covers around 30% of facial skin.

The maxillary division V2 gives rise to six sensory branches, of which two are responsible for the sensory innervation of the hard palate inside the oral cavity (greater palatine and nasopalatine nerves). The mandibular division V3 includes five sensory branches, of which the lingual nerve and the buccal nerve innervate the floor of the oral cavity and the inside of the cheeks. Thus, four out of 11 branches of the V2 and V3 divisions innervate the inside of the mouth. Combining this fact with recent histological analyses, which found that skin within the V3 innervation area contains almost twice the number of fibers than skin innervated by V1 (68), suggests by rough approximation that around 25-30% of the 62,0000 myelinated fibers of the trigeminal nerve are responsible for the sensory innervation of the oral mucosa, leaving around 43,000 to innervate the facial skin and lips.

The pattern of the sensory innervation changes across the three divisions and the density of myelinated fibers was estimated by Nolano et al. (68) at 8.0, 15.9, and 16.4 mm^2^ in V1, V2, and V3, respectively. These estimates include multiple branches originating from the same afferent and also count any fibers merely traversing a given skin area rather than terminating there, and thus cannot be used directly to estimate the number of individual afferents. Nevertheless, in relative terms, these histological counts can be expected to scale similarly to the actual afferent counts. To arrive at estimated innervation densities for V1, V2, and V3, we, therefore, divided the total number of fibers estimated above across V1, V2, and V3 in the proportions estimated by Nolano et al. (68). A total facial skin area of 675 cm^2^ was assumed (69).

### Hairy skin on arms, trunk, and legs

The dorsal roots of the spinal cord contain 1-1.2 million fibers in total, ranging from large, medium, and small myelinated to unmyelinated nerve fibers (70, 71). The region of skin innervated by all tactile afferents passing through a given dorsal root is known as a dermatome. While the specific territory innervated by each dermatome varies between people, and dermatomes also generally overlap within individuals, they nevertheless follow a systematic pattern. Fiber counts in individual dorsal roots can, therefore, be used to estimate the innervation of their associated dermatomes. For the estimates presented in this study, we follow the recent fiber counts in Liu et al. (70). The territory of each dermatome is derived from illustrations in Grant and Anderson (72): the area of skin innervated by each dermatome is calculated as the sum of the areas covering the front and back of the body, respectively, assuming a total area of skin of 1.5 m^2^ (73).

Only a subset of dorsal root fibers will be myelinated fibers in the Aβ range and underlie tactile innervation of the skin, rather than internal organs. The fraction of myelinated fibers varies across the spinal tract, being higher in the cervical and lumbar tract (74) and, on average, around 40% of axons have been classified as unmyelinated (75). Assuming a 50:50 split into Aδ and Aβ of the myelinated fibers, and considering a further 10% of Aβ fibers innervating deep structures (66), an average fraction of 16% (range: 10%-25%) of the fibers in the dorsal roots are estimated to be Aβ fibers involved in the transmission of tactile sensations. In the dermatomes C6-C8, for example, we estimate that about 53,500 of the 210,000 fibers are tactile afferents. Considering that approximately 36,000 of these are in the ulnar and median nerve and innervate the glabrous skin of the hand, the remaining tactile afferents in the C6-C8 dermatomes cover the hairy skin of the hand dorsum and forearm. This calculation leads to an estimated density of 9.4 units/cm in these areas, which is consistent with the overall hairy skin estimations here proposed. Our estimates also agree well with a recent count in the L4 and L5 dorsal roots that found around 30% of axons had a diameter bigger than 5 μm, not all of which contribute to the tactile innervation of the skin (76).

The main source of uncertainty in our estimates is the total number of fibers in the dorsal root and, most importantly, the proportion of myelinated Aβ fibers for each dermatome (74, 76). This question has only been investigated experimentally in a subset of dorsal roots, and different studies report conflicting results. For this reason, we repeated our calculations assuming a possible positive or negative variation of 20% on the number of tactile afferents for each dermatome. When doing so, we noticed that we fell short of or exceeded the physiologically plausible range in several instances, lending credibility to our original estimates. For example, in dermatomes C6, C7, and C8 considering that 18,000 of the total number of afferents can be expected to innervate the palmar surface of the hand, a reduction of more than 20% of the dorsal root fiber portion considered as tactile afferents, would result in a number close to zero (or even negative) for tactile afferents innervating the hairy skin of the back of the hand and part of the forearm. Similarly, an increase of more than 20% in the portion of dorsal root fibers considered to be tactile afferents would, in some dermatomes such as L1, result in areas of hairy skin having a density of afferents improbably close to that of the hairless skin of the foot and some areas of the palm of the hand. Taking these limits into account, the overall number of tactile afferents innervating the hairy skin is likely to fall in the range 110,000-180,000, leading to a total number of 200,000-270,00 afferents across the whole body.

### Visualisation

Figures showing body regions scaled by their innervation density (‘homunculi’) were generated using a flow-based algorithm from 2D body region outlines (77).

## ACKNOWLEDGEMENTS

We would like to thank Roland Johansson, Ingvars Birznieks, Laura Edmondson, and Rodrigo Kazu Siqueira for helpful comments on the manuscript, as well as Tom Farrow for neuroanatomical guidance. This work was supported by the Wellcome Trust [209998/Z/17/Z] and by the EU Horizon 2020 research and innovation programme under grant agreement 813713 (NeuTouch).

